# Calcite seed-assisted microbial induced carbonate precipitation (MICP) and its potential in biocementation

**DOI:** 10.1101/2020.07.17.206516

**Authors:** Jennifer Zehner, Anja Røyne, Pawel Sikorski

## Abstract

Microbial-induced calcium carbonate precipitation (MICP) is a biological process inducing biomineralization of CaCO_3_. This can be used to form a solid, concrete-like material. To be able to use MICP successfully for producing solid materials, it is important to understand the formation process of the material in detail. It is well known, that crystallization surfaces can influence the precipitation process. Therefore, we present in this contribution a systematic study investigating the influence of calcite seeds on the MICP processes. We focus on the pH changes during the crystallization process measured with absorption spectroscopy and on the optical density (OD) signal to analyze the precipitation process. Furthermore, optical microscopy was used to visualize the precipitation processes in the sample and connect them to changes in pH and OD. We show that there is a significant difference in the pH evolution between samples with and without calcite seeds present and that the shape of the pH evolution and the changes in OD can give detailed information about the mineral precipitation and transformations. In the presented experiments we show that amorphous calcium carbonate (ACC) can also precipitate in the presence of initial calcite seeds, which can have consequences for consolidated MICP materials.

## 1 Introduction

Biocementation is a biological process which increases the engineering properties of a granular medium by biomineralization. Biomineralized CaCO_3_ precipitates fill pores and bind the granular medium together, which results in an increase of strength and stiffness, and reduce the permeability of the granular medium^1^. One of the most commonly used process to achieve biocementation is microbially induced calcite precipitation (MICP). MICP has been successfully used for soil stabilization^1,2^, which reduces the environmental impact of ground improvement, since conventional ground improvement techniques often have a high impact on the environment^3^. Moreover, MICP has the potential to replace conventional concrete as a building material in some applications^4^. Conventional concrete production contributes with up to 5% to the anthropogenic CO_2_ emissions^5^, and therefore there is a need for CO_2_ reduction within the construction industry. MICP based solid materials have the potential to lower the CO_2_ emissions in concrete production^6^. MICP is commonly based on the urea hydrolysis reaction catalyzed by the urease enzyme^7,8^:

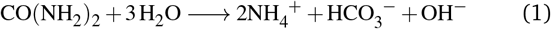

Urea hydrolysis leads to the production of bicarbonate and a pH increase. In the presence of sufficient concentration of calcium ions this can result in CaCO_3_ precipitation:

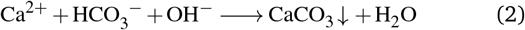

Supersaturation is the driving force for precipitation, which can take place once the saturation state in the system *S* > 1^9^:

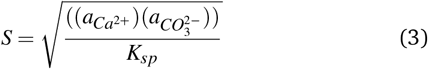

where 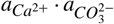 is the ion activity product of calcium and carbonate ions in the solution, and K_*sp*_ is the solubility product of the nucleating polymorph of CaCO_3_. A sufficient level of supersaturation results in nucleation and is followed by subsequent crystal growth. In MICP processes the supersaturation is controlled by the pH value, the amount of dissolved inorganic carbon and the calcium concentration^8^.

In MICP, several injections of crystallization solution are typically used to achieve sufficiently consolidated granular material^10–12^. During initial injections calcite crystals will nucleate within the granular medium. Therefore, not only the granular medium, the bacteria cells, and gas bubbles in the system, but also calcite crystals need to be considered in the material forming process. The presence of such surfaces in the crystallization solution results in heterogeneous nucleation. In heterogeneous nucleation the energy barrier necessary for nucleation is lowered and the nucleation rate increases^13^.

The suitability of different surfaces as nucleation sites in heterogeneous nucleation has been investigated previously. Lioliou *et al*.^14^ compared the nucleation of CaCO_3_ in the presence of calcite and quartz seeds, and found that nucleation can take place at lower supersaturation levels for calcite seeds compared with quartz seeds. It was also shown by Dawe *et al*. that gas bubbles in the crystallization system promote CaCO_3_ precipitation^15^. Furthermore, the influence of bacteria cell surfaces as nucleation sites for MICP has been investigated. While Mitchell *et al*. showed that bacteria surfaces are not good nucleation sites^16^, it has been shown recently by Ghosh *et al*. that nanoscale calcium carbonate crystals can form on the surface of *Sporosarcina pasteurii*^17^. *S. pasteurii* is an extensively studied microorganism for MICP processes ^7,18,19^.

We have previously reported an experimental method using absorption spectroscopy to study the precipitation process and the real time pH evolution of MICP without calcite seeds in volumes sufficiently small that mixing and transport occurred only by diffusion^20^. In this crystallization process, where there were no initial calcite crystals present, three stages could be identified. In the first stage, urea hydrolysis catalyzed by bacterial urease increased the pH, and during this phase precipitation of metastable amorphous calcium carbonate (ACC) and vaterite was observed (Stage I). After a short stable pH phase nucleation of calcite was observed (Stage II). Nucleation and growth of calcite crystals resulted in a pH decrease (Stage III), while the metastable ACC/vaterite was observed to dissolve.

To better understand the formation of an MICP consolidated material it is important to know how the presence of calcite crystals will influence the crystallization. Therefore, we have investigated the influence of calcite seeds on the crystallization process in MICP. We compare the precipitation in samples with and without calcite seeds and investigate the precipitation kinetics by monitoring the pH evolution in small volumes with absorption spectroscopy. From the changes in the optical density of the reaction solution, we determine the onset of the precipitation and follow phase transformations. We show that the pH evolution is significantly different between unseeded and seeded samples and can give information about the precipitation process. Furthermore, we demonstrate that precipitation of ACC and vaterite can also occur for samples with calcite seeds present and that this process depends on the bacteria concentration and consequently on the urease activity.

## 2 Material and methods

All solutions were filtered with a polycarbonate (0.22 *μ*m) syringe filter before use. The used chemicals were purchased at Sigma-Aldich (Norway), unless otherwise stated. For preparation of the crystallization solution as well as for the bacteria culture, the same protocol as in previously presented work has been used^20^, and a short description of the used protocols is given below.

### 2.1 Crystallization solution

The crystallization solution for the presented experiments consisted out of two components: urea, and dissolved chalk solution. The initial concentration of urea before the start of the hydrolysis reaction was 0.1 M. Dissolved chalk solution (DCS) was used as a calcium source for MICP experiments. Crushed limestone (Franzefoss Miljøkalk AS (Norway)) was dissolved with 300mM lactic acid. After 24 h reaction time the undissolved parts of the crushed limestone were filtered out with a 0.22 *μ*m syringe filter. Urease producing bacteria cells were added to the crystallization solution to catalyze the hydrolysis reaction. The amount of the bacteria dilution contributed with 10% to the final volume of the sample.

### 2.2 Bacteria culture

For the presented experiments the urease-producing bacterium *Sporosarcina pasteurii* (Strain DSM33 of *S. pasteurii*) purchased from “Deutsche Sammlung von Mikroorganismen and Zellkulturen” (DSMZ) was used. DSMZ medium 220 with pH of 7.3 supplemented with urea and consisting of 15 gl^−1^ peptone from casein, 5gl^−1^ peptone from soy meal, 5gl^−1^ NaCl and 20gl^−1^ urea was used as growth medium. The culture was inoculated with 1% frozen glycerol stock. After inoculation the culture was incubated overnight (17 h) at 30 °C with constant shaking. Afterward the culture was incubated without shaking at 30 °C until further use 24 h after inoculation. The cultures were prepared for the experiment by centrifuging the culture at 4800 ×*g* for 8 min and the cells were subsequently washed twice with pre-warmed 0.01 M PBS (phosphate buffered saline) to remove the growth medium. The cells were re-suspended and diluted with 0.01 M PBS. Three different bacteria cultures were used for the presented experiments. As an indication of bacterial cell concentration the optical density (OD_600nm_) at 600 nm was measured in a 96-well plate (volume: 150 *μ*m) before adding the bacteria suspension to the crystallization solution. The optical densities for the bacterial suspensions used in the presented experiments as well as the used sample names are shown in Table 1.

**Table 1.** Bacteria cell concentrations of the cultures added to the crystallization solution. The table shows the name and the OD_600nm_ of the bacteria dilutions, which were added to the crystallization solution. The ratio of bacteria dilution to sample volume was 1 to 10. Additionally, the table gives an overview of the sample names, which were used for unseeded and seeded sample where the bacteria dilutions have been added.

| Dilution name | OD<sub>600nm</sub> | Bacteria culture | sample name |  |
| --- | --- | --- | --- | --- |
| pH monitoring |  |  | unseeded | seeded |
| D1-C1 | 1.40 | culture1 | D1-C1<sub>us</sub> | D1-C1<sub>s</sub> |
| D2-C1 | 1.30 | culture1 | D2-C1<sub>us</sub> | D2-C1<sub>s</sub> |
| D3-C1 | 1.11 | culture1 | D3-C1<sub>us</sub> | D3-C1<sub>s</sub> |
| D4-C1 | 0.64 | culture1 | D4-C1<sub>us</sub> | D4-C1<sub>s</sub> |
| Optical microscope |  |  | unseeded | seeded |
| D2-C2 | 0.88 | culture2 | — | D2-C2<sub>s</sub> |
| D1-C3 | 0.89 | culture3 | D1-C3<sub>us</sub> | — |
| Confocal laser scanning microscope |  |  | unseeded | seeded |
| D1-C2 | 0.96 | culture2 | — | D1-C2<sub>s</sub> |

The dilution of the original cultures with PBS is shown in Table S1, Supplementary Information. The optical density values of bacteria cultures given in Table 1 refer to the OD_600nm_ of the bacteria culture before adding the bacteria cells to the crystallization solution.

### 2.3 Calcite seeds

Calcite seeds for optical microscopy and global pH monitoring experiments were prepared by mixing 0.2 M CaCl_2_ solution with 0.2 M Na_2_CO_3_ solution in an airtight reactor. The crystallization reaction was performed at 10 °C and the solution was stirred with a mechanical stirrer for 48 h. After the reaction was completed, the calcite seeds were washed with water and ethanol before further use. The finished calcite seeds have been characterized with scanning electron microscopy. The calcite seeds had a rhombohedral shape with a width of about 8 *μ*m (see Supplementary Information: Figure S1).

Additionally, for confocal laser scanning microscopy experiments a fluorescent dye was incorporated in the calcite seeds during the crystal growth, following a procedure reported by Green *et al*. ^21^. The fluorescent calcite seeds were directly grown on glass microscope cover-slides, as follows: The slides were placed in a beaker with CaCl_2_ solution and 0.1 mM solution of the fluorescent dye HPTS (8-Hydroxypyrene-1,3,6-trisulfonic acid). To initiate the crystallization, Na_2_CO_3_ solution was added and mixed thoroughly. The concentration of Ca^2+^ and CO_3_^2-^ was 5mM. The reaction was left to proceed for 3 days. Afterwards, the slides were washed with DI water and ethanol before use. The shape of the crystals with fluorescent dye incorporated are shown in Figure S2, Supplementary Information.

### 2.4 Real time pH monitoring

Real time pH evolution measurements were performed with a method reported earlier to measure the average pH in small volumes (200 *μ*L)^20^. Crystallization solution and the pH indicator Phenol Red (0.4 *μ*M) were pre-mixed in a 96-well plate, and bacteria suspensions were added to start the reaction. To minimize the gas exchange with the environment the well plate was covered with a transparent tape before the measurement was started. The change in the optical density (OD) at 558 nm and 750 nm was measured with a spectrophotometer (SpectraMax^®^ i3 Platform). No signal from the pH sensitive dye was detected above 600 nm, meaning that the change in OD at 750 nm (OD_750nm_) was only caused by scattering of precipitates, seeds and bacteria cells in the sample. Consequently, the OD_750nm_ could be used for background corrections for the OD at 558 nm. The background corrected OD at 558 nm corresponds to the absorbance of the sample, and the absorbance was used to calculate the pH, using a standard-curve created by recording the background corrected OD for calibration buffers with known pH. The OD_750nm_ was also used to monitor changes in amount of CaCO_3_ in the sample, since the change in OD_750nm_ due to bacterial growth is expected to be small compared with the change due to precipitation of CaCO_3_.

### 2.5 Optical microscopy

The crystallization process with and without calcite seeds was monitored with an optical microscope (Motic, AE31E). The crystallization reaction was performed in a 96-well plate (sample volume: 200 *μ*L). The objectives used were 10x (0.25 NA) for crystallization without seeds, and 20x (0.3 NA) for crystallization with seeds. Higher magnification was used for the seeded samples due to the small size of the calcite seeds. Images were recorded with a Moticam 5.0.

### 2.6 Confocal laser scanning microscopy

A confocal laser scanning microscope (CLSM, Leica TCS SP5) using an Argon laser (488 nm) and detection range 500 nm to 550 nm was used to investigate the crystal growth in the presence of fluorescent calcite seeds described above, and to distinguish the initial calcite crystals and the microbial-induced CaCO_3_ precipitation.

### 2.7 Scanning electron microscopy

Scanning electron microscopy (SEM, Hitachi S-3400N) was used for characterizing the precipitated crystals. For SEM characterization the crystals were first dried on filter paper and the dry crystals were attached to carbon tape on the SEM stub. The crystals were sputter coated with a 10 nm layer of Pt/Pd (80/20) using Cressington 208 HR sputter coater.

## 3 Results

### 3.1 Unseeded experiments

The pH evolution as well as the OD_750nm_ curves for CaCO_3_ precipitation induced by hydrolysis of urea by *S. pasteurii* in samples that did not contain calcite seeds are shown in Figure 1a,b. Experiments were performed using four different bacteria cell concentrations as specified in Table 1.

**Fig. 1.**
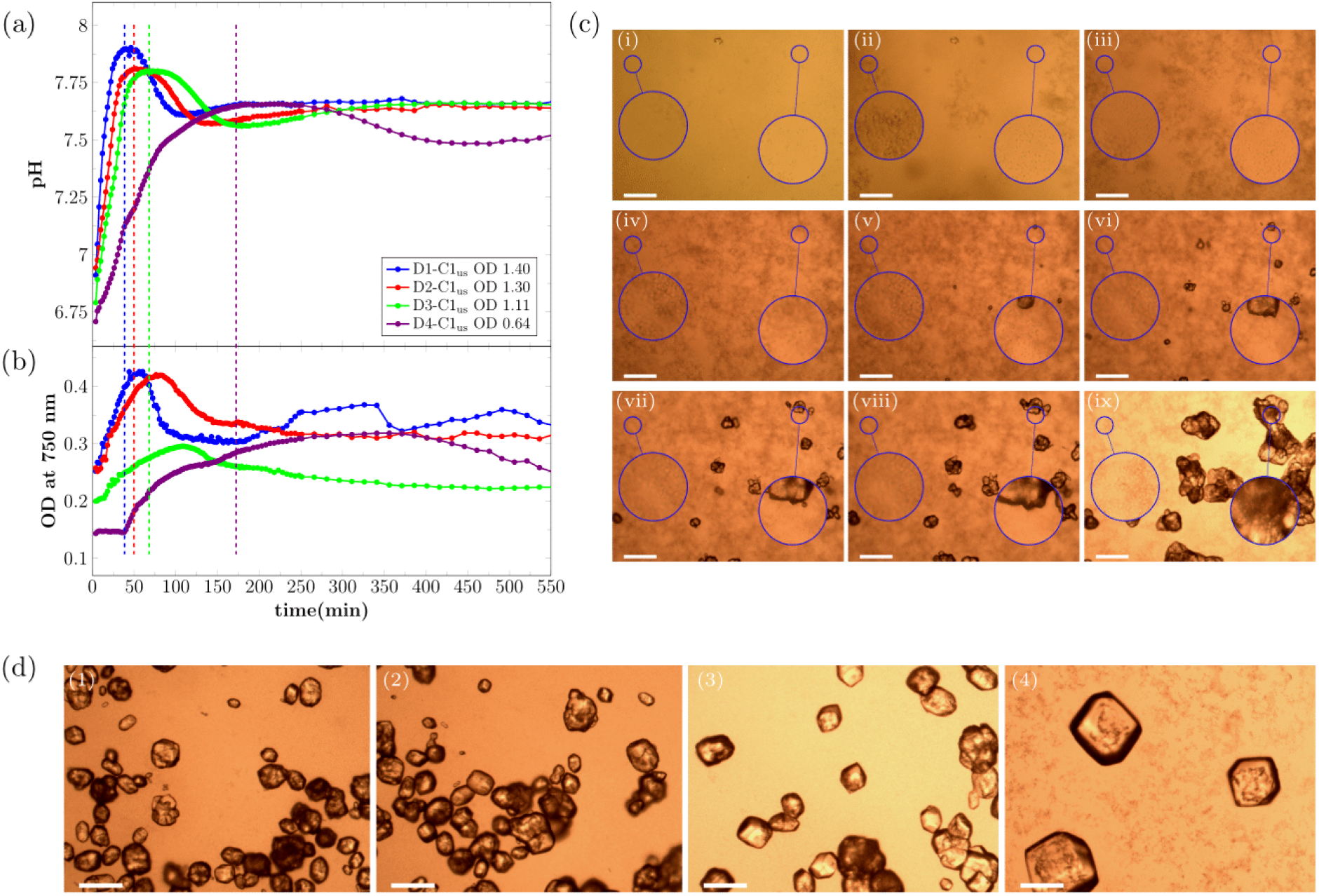
Real time evolution of (a) pH and (b) OD_750nm_ for MICP experiments without the presence of calcite seeds for the first 550 min of the reaction. Note that the first shown data point of the measurement is taken 4 min after the start of the reaction (= 0 min). The signal of the OD_750nm_ correlates with the amount of CaCO_3_ in the sample. Dashed vertical lines mark the time-point of maximum pH. The OD in the legend shows the OD_600nm_ of the bacteria dilution before adding to the crystallization solution. (c) Optical microscope time-series of precipitation process for bacteria dilution D1-C3 (OD_600nm_ =0.89) without calcite seeds present for different time-points of the reaction: (i) 0 min, (ii) 14 min, (iii) 23 min, (iv) 47 min, (v) 54 min, (vi) 88 min, (vii) 148 min, (viii) 240 min, and (ix) 21 h. The magnified area on the left shows an area where no calcite will nucleate during the experiment. The magnified area on the right shows an area where a calcite crystal will nucleate. (d) Micrographs of precipitated crystals after 21 h reaction time for (1) D1-C1_us_, (2) D2-C1_us_, (3) D3-C1_us_, and (4) D4-C1_us_ without the presence of calcite seeds. The scale-bar in (c) and (d) is 150 *μ*m.

As expected, the rate of the pH increase at the beginning of the reaction was correlated with the bacterial cell concentration and therefore with the urea hydrolysis rate (Figure 1a). The highest pH was reached for the samples that contained the highest bacterial cell concentration. After a short period of stable pH, the pH decreased and this trend was observed for all four tested bacterial cell concentrations. The pH remained stable longer for lower bacterial cell concentrations. Furthermore, it could be observed that the pH increased in the final stage of the reaction before stabilizing.

The observed changes in the pH of the crystallization solution were correlated with changes in OD_750nm_, which measures how much light is scattered by the sample. For the two highest bacteria cell concentrations, the OD_750nm_ increased fast from the beginning of the reaction and decreased after the maximum pH was reached in the sample. For the two lowest bacteria cell concentrations, the OD_750nm_ was nearly stable at the beginning of the reaction, before increasing after 14 min and 38 min for D3-C1_us_ and D4-C1_us_, respectively.

The increase in the OD_750nm_ during the first phase of the reaction can be interpreted as precipitation of amorphous calcium carbonate (ACC), which quickly transformed into vaterite ^20^. This was followed by the nucleation of calcite, accompanied by the dissolution of ACC/vaterite. The dissolution of the metastable phase resulted in a decrease of the OD_750nm_ for all four bacteria cell concentrations. This is a result of fewer, larger crystals scattering less light than many, small particles. The pH of the sample started to decrease once calcite nucleated (See discussion and Figure 5 for more detailed description of the various processes taking place during precipitation). The second period of pH increase started at the same time as the OD_750nm_ signal started to stabilize. This second period of pH increase was most likely caused by a slowing down of the calcite precipitation rate due to a low amount of calcium remaining in solution.

This interpretation of the changes in pH and OD_750nm_ is supported by optical microscopy. Figure 1c shows selected timepoints for the precipitation reaction for the sample D1-C3_us_ (see Table 1). The precipitation of small particles was observed in the initial stage of the reaction (Figure 1c, i-iv). These precipitates have been characterized previously^20^ and identified as ACC, which quickly transforms into vaterite. After the nucleation of calcite crystals (Figure 1c, v), the calcite crystals grew and the ACC/vaterite phase dissolved. The dissolution was fastest near existing calcite crystals (Figure 1c, v-viii, magnified area on the right).

Figure 1d shows crystals precipitated in the experiments described above (Figure 1a,b) imaged after the reaction was completed (21 h). It was observed that the bacteria cell concentration had a significant effect on the morphology of the precipitated crystals. For the highest bacteria cell concentration (D1-C1) the smallest crystals, with a mixture of rhombohedral and spherical morphology were observed (Figure 1d, 1). For samples with lower bacteria cell concentration, the precipitated crystals were larger and had a more regular, rhombohedral shape (Figure 1d, 2-4)).

### 3.2 Seeded experiments

The presence of calcite seeds had two effects on the MICP experiments (see Figure 2a,b). First, the seeds affected the pH of the crystallization solution before the onset of the urea hydrolysis reaction, due to partial dissolution of the calcite seeds in the initially under-saturated solution. The starting pH was around 5.3 for unseeded and 6.1 for seeded samples (see Table S2, Supplementary Information). Secondly, seeds affected the precipitation process as the calcite seeds acted as nucleation sites for heterogeneous nucleation.

As for the unseeded samples, the pH increase at the beginning of the reaction was correlated to the bacterial cell concentration (Figure 2a). The period of rapid pH increase was followed by a pH decrease for the three highest bacteria cell concentrations. For the lowest bacteria cell concentration, no pH decrease was observed, and the pH stabilized after the initial pH increase.

**Fig. 2.**
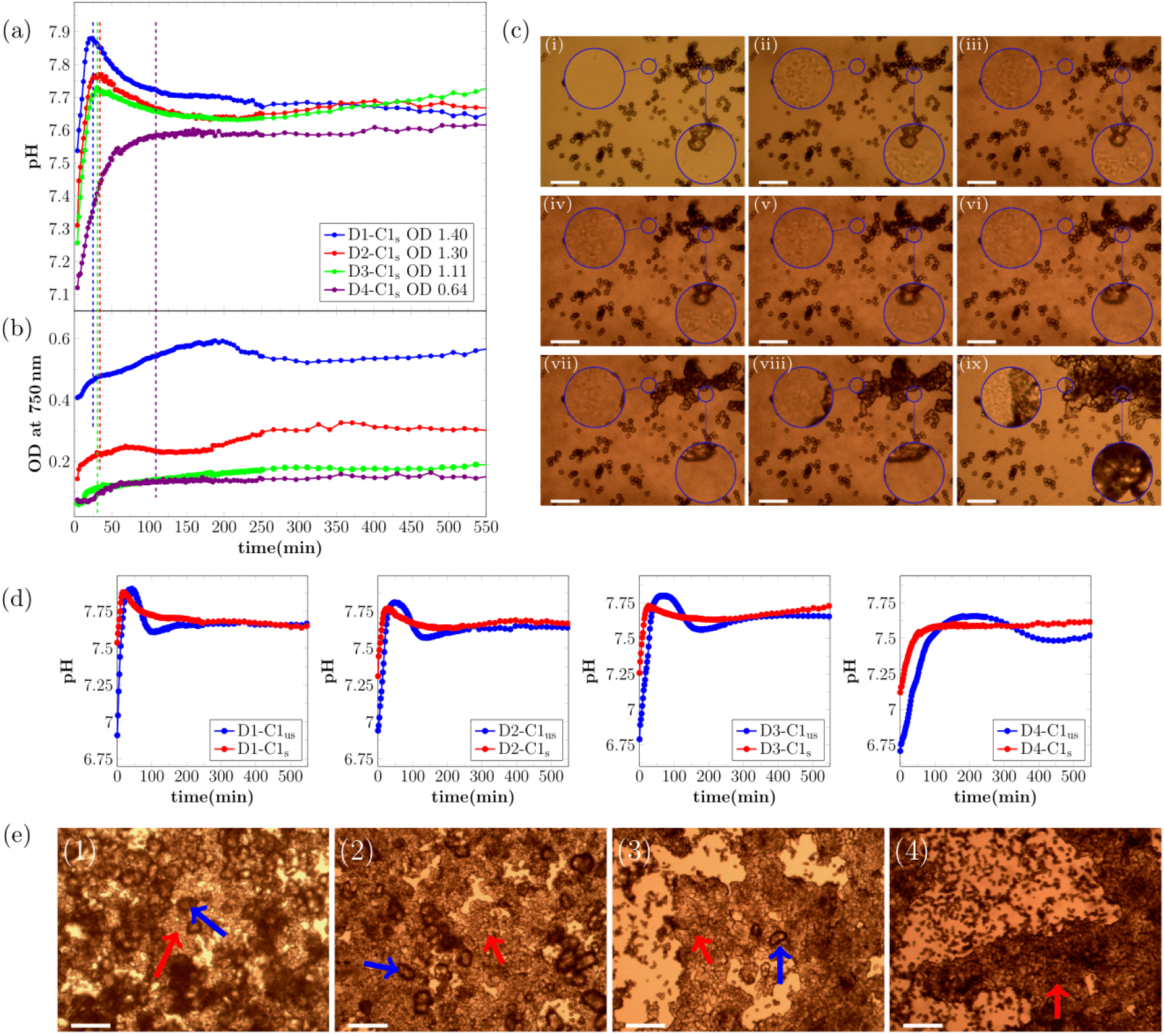
Real-time evolution of (a) pH and (b) OD_750nm_ for MICP experiments with the presence of calcite seeds. Note that the first shown data point of the measurement is taken 4 min after the start of the reaction. The OD_750nm_ was measured before start of the reaction to get the contribution of the initial seeds in the crystallization solution, and this starting value was subtracted form the OD_750nm_. Therefore, the shown signal is correlated to the amount of precipitated CaCO_3_. The OD in the legend shows the OD_600nm_ of the bacteria dilution before adding to the crystallization solution. (c) Optical microscope time-series of the precipitation process for bacteria dilution with OD_600nm_=0.88 with calcite seeds present for different time-points of the reaction: (i) 0 min, (ii) 20 min, (iii) 28 min, (iv) 50 min, (v) 80 min, (vi) 110 min, (vii) 170 min, (viii) 230 min, and (ix) 21 h. The magnified area on the right shows calcite seeds, which will increase in size during the experiment. The magnified area on the left shows an area with a small distance to calcite seeds which will increase in size during the experiment. (d) Comparison of real time pH evolution of MICP experiments with and without calcite seeds. (e) Micrographs of precipitated crystals after 21 h reaction time for (1) D1-C1_s_, (2) D2-C1_s_, (3) D3-C1_s_, and (4) D4-C1_s_. The red arrows indicate calcite seeds which grew during the experiment and the blue arrows indicate additional nucleated calcite crystals. The scale-bar is 50 *μ*m in (c) and 150 *μ*m in (e).

As seen in Figure 2d, the shape of the pH curves is significantly different for unseeded and seeded samples. The initial pH increase had approximately the same slope for both unseeded and seeded samples at the start of the reaction. The offset along the time axis between unseeded and seeded pH curves was caused by the higher starting pH of the seeded samples (see Table S2, Supplementary Information). The maximum pH was lower for the seeded samples. This was probably caused by an onset of nucleation and crystal growth at lower supersaturation levels due to the existing calcite surfaces. For unseeded samples a stable pH phase at the maximum pH was observed, which was longer for lower bacterial cell concentrations. For the three highest bacteria cell concentrations, the pH decrease was faster for unseeded than for seeded samples. For the lowest bacterial cell concentration no pH decrease was observed in the presence of calcite seeds (Figure 2d).

The OD_750nm_ of the seeded samples was measured before adding the bacteria cells, and this value was subtracted from the subsequent real time OD_750nm_ measurement during the precipitation process, to ensure the OD_750nm_ signal shown in Figure 1b does not include the contribution of the calcite seeds. Changes in the sample OD_750nm_ in Figure 1b were therefore mainly caused by the precipitation of CaCO_3_. For the two highest bacteria cell concentrations, the OD_750nm_ of the samples increased throughout the reaction. For the two lowest bacteria cell concentrations, the OD_750nm_ of the sample was constant at the beginning of the reaction and started to increase after 14 min and 28 min for D3-C1_s_ and D4-C1_s_, respectively. The increase in OD_750nm_ was observed for both samples only during the pH increase in the initial phase of the reaction.

The precipitation process for the sample D2-C2 (see Table 1) was investigated with optical microscopy (Figure 2c) to correlate changes in the pH and the OD_750nm_ to the precipitation process. Similar to the unseeded experiments, precipitation of small particles was observed in the initial phase of the reaction (Figure 2c, ii-iii). The morphology of the precipitated particles was similar to what was observed in the unseeded experiments and therefore the initial precipitation could be identified as ACC/vaterite. The precipitation of ACC/vaterite was also indicated by a change in the OD_750nm_ of the samples during the initial reaction stage. The initial precipitation of ACC/vaterite was followed by growth of the crystal seeds present in the sample (Figure 2c, iv-ix), and the ACC/vaterite dissolved at the later stage of the reaction (see magnified areas in Figure 2c, v-vii) with faster dissolution in close proximity of the growing crystals (magnified area on the right). The dissolved ACC/vaterite reprecipitated on the surface of the seeds, but no pronounced decrease in the OD_750nm_ was observed, most likely due to the fact that less calcium was bound in metastable precipitates in seeded samples.

For the two lowest bacteria cell concentrations, the onset of precipitation occurred earlier in the seeded than in the unseeded experiments. This is likely a consequence of the higher starting pH of the crystallization solution, due to partly dissolved calcite seeds (Table S2, Supplementary Information).

Optical micrographs of crystals precipitated in seeded experiments recorded after 21 h are shown in Figure 2e. For a low hydrolysis rate, only the growth on calcite seeds was observed (red arrow in Figure 2e, 4). However, for the three highest bacteria cell concentrations, which corresponded to a higher hydrolysis rate, additional nucleated calcite crystals were observed in the samples. The additional nucleated crystals are marked with a blue arrow and calcite seeds which increased in size during the reaction are marked with a red arrow (Figure 2e, 1-3). The additional precipitated calcite crystals in Figure 2e had a more irregular shape compared to samples without calcite seeds present Figure 2.

The growth process on the surface of the calcite seeds was investigated in more detail with confocal laser scanning microscopy (Figure 3a-f). The growth process is shown for six time-points. Figure 3a shows the calcite seed 1 min before the crystal growth started. The crystal growth on the surface of the calcite seed started with small islands on the calcite crystal (Figure 3b) and the islands increased in size over the 13 min (Figure 3c-e), until the growing crystals formed a new layer of calcite on the surface of the calcite seed (Figure 3f). The incorporation of a fluorescent dye into the initial calcite seed crystal (see Materials & Methods) made it possible to distinguish between the initial crystal seeds (originated from an abiotic process) and the newly grown calcite crystals (originated from a biotic) on the seed surface (Figure 3g).

**Fig. 3.**
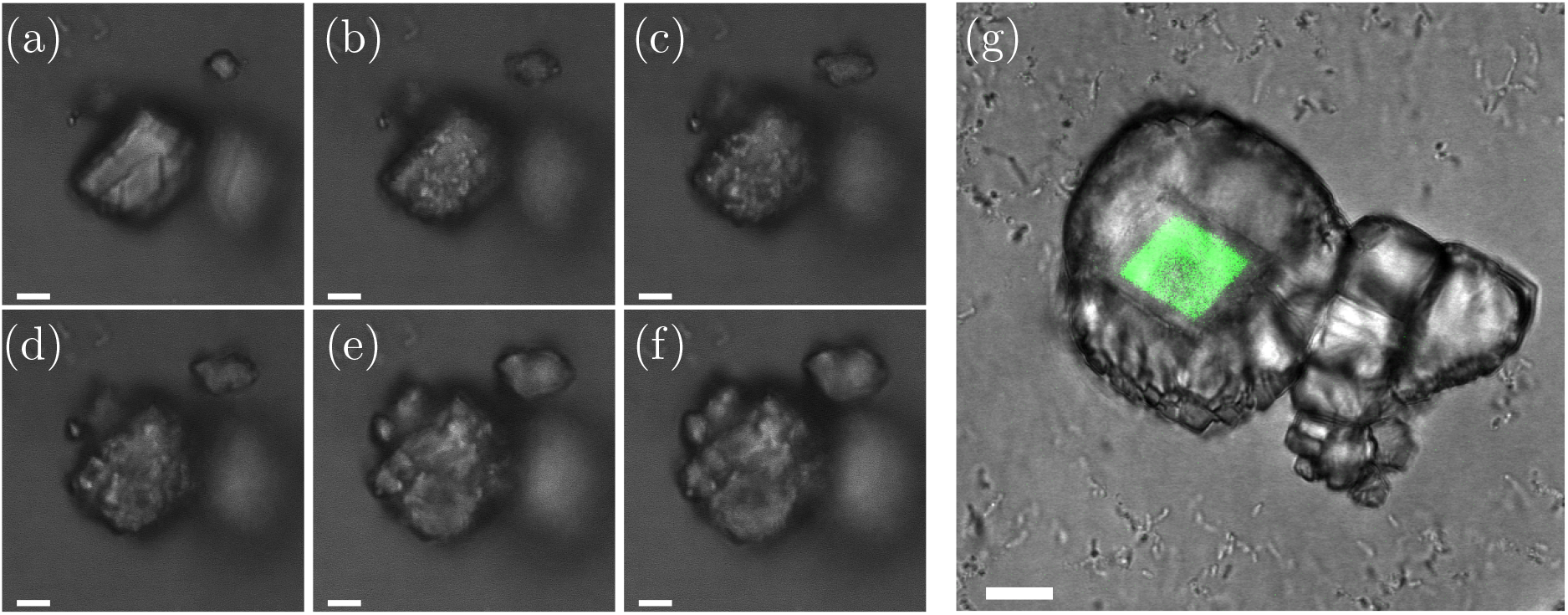
(a-f) Growth process of calcite seeds. Images are taken with a CLSM. Images are taken (a) 1 min before the growth process started, and (b) 1 min (c) 3 min (d) 6 min, (e) 11 min and (f) 14 min after the growth process started. (g) Finished crystal. The original fluorescent calcite seed (originated from an abiotic process) had a fluorescent dye incorporated and was visualized with CLSM. The scale-bar is 10 *μ*m

After the reaction was finished (after 2 d) the precipitated crystals shown in Figure 1d and Figure 2e were removed from the wells and transferred to filter paper. Dried crystals for unseeded and seeded samples for two bacterial cell concentrations were characterized with SEM (Figure 4). For the unseeded samples it can be seen that the precipitated calcite crystals are bigger for the lower bacterial cell concentration (Figure 4a,c). Furthermore, for D1-C1 the imprints of bacterial cells as well as bacterial cell can be identified on the crystal surface (small insert in Figure 4a). Due to the lower bacterial cell concentration less bacterial imprints and bacterial cells could be detected on the surface of the calcite crystal of D3-C1 resulting in a smoother crystal surface (Figure 4c). Small, needle shape crystals on the surface of the crystal are most likely calcium lactate crystals, since the crystals were not washed before the drying and characterization process. Furthermore, the bottom side of the crystals was analyzed for the unseeded samples (Figure 4b,d). The bottom-side of the crystals is the part of the crystals which was growing on the well surface. Here we could detect, that for the highest bacterial cell concentration (Figure 4b), the amorphous phases was encapsulated by the growing crystal. This was most likely caused by the fact that a fast growth process isolated some amount of the initially precipitated amorphous phase from the crystallization solution. Since the transformation process of the amorphous phases to calcite is a dissolution based transformation, the transformation of the amorphous phases was suppressed. No amorphous phases were detected on the bottom of the crystal for the lower bacterial cell concentration (Figure 4d). SEM images of ACC have been reported previously ^22,23^ and are in good agreement with the morphology of the amorphous phase shown in Figure 4b. For the seeded samples the initial calcite seeds as well as seeds which grew during the experiments could be observed in the SEM micrographs (Figure 4e-h). For D1-C1 seed crystals which grew during the experiment could not be identified due to precipitation at high supersaturation (Figure 4e,f). For the lower bacterial cell concentration the initial calcite seeds (originated from an abiotic process) and the precipitated crystals (originated from MICP process) could be distinguished (Figure 4g,h). It can be seen that the precipitated crystals grew on the surface of the calcite seeds, and connecting them together (Figure 4g). Furthermore, initial calcite seeds have a different surface compared to the MICP induced calcite, as can be seen in Figure 4h.

**Fig. 4.**
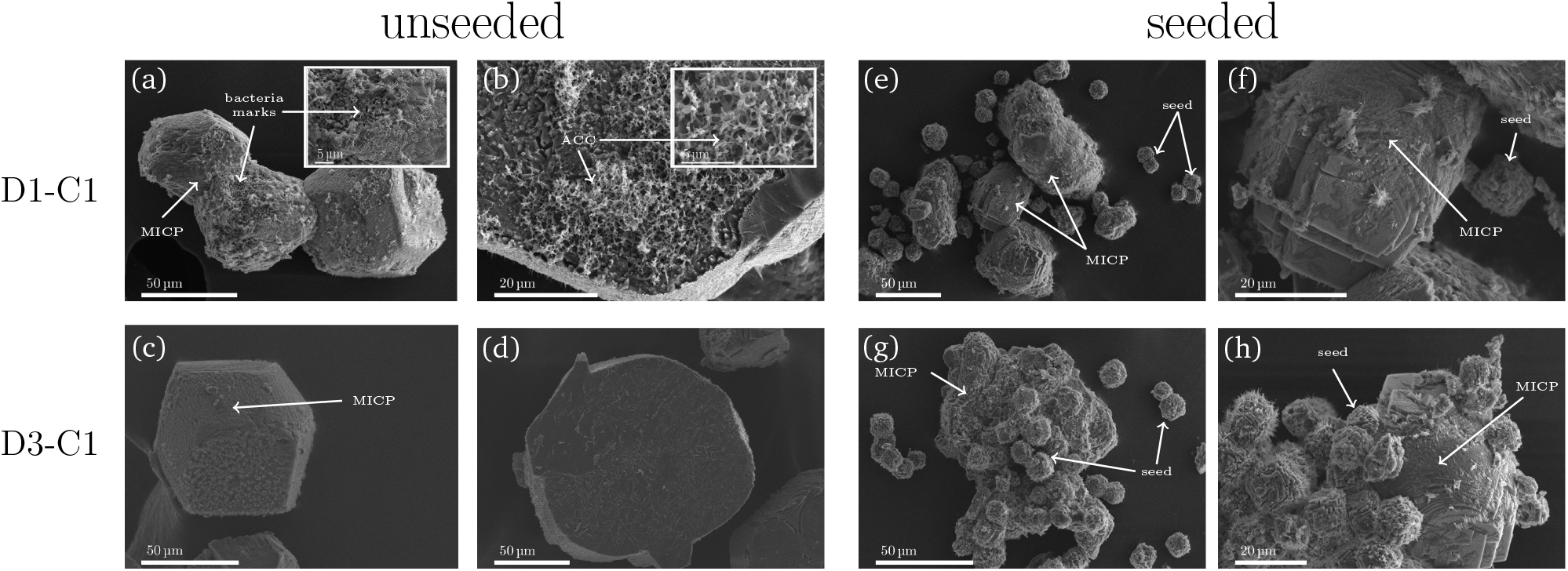
SEM characterization of precipitated crystals for unseeded and seeded samples for high bacterial concentration (D1-C1) and lower bacterial cell concentration (D3-C1). CaCO_3_ crystals which originated from MICP processes are marked with MICP in the SEM micrographs. (a) MICP induced calcite crystals for D1-C1 in unseeded experiments. Imprints of bacteria can be identified on the surface. (b) Bottom of the crystal for D1-C1 (unseeded) were the amorphous phase is enclosed in the calcite crystal. (c) MICP induced calcite crystals for D3-C1 in unseeded experiments and (d) the bottom of the crystals for D3-C1 in unseeded experiments, where no amorphous phase was detected. (e,f) Seeded samples for D1-C1. Initial calcite seeds (which did not grow during the experiment) and the MICP induced CaCO_3_ can be seen. (g,h) Initial calcite seeds (which did not grew during the experiment) and the precipitated CaCO_3_ can be seen. The initial calcite seeds can be identified within the precipitated CaCO_3_. Before SEM characterization, the crystals were transferred from the well plate to filter paper for drying. The crystals were not washed before SEM characterization. Needle-like crystals on the calcite surfaces are most likely calcium lactate.

## 4 Discussion

In the presented experiments we investigated the influence of calcite seeds on the CaCO_3_ precipitation in MICP processes.

We observe significant differences in the pH evolution for unseeded and seeded samples (Figure 2d). These differences can be connected to the saturation state in the sample and consequential to different processes during the precipitation. A schematic illustration of the connection of the saturation state to the pH evolution in unseeded and seeded samples with high bacterial cell concentration is shown in Figure 5.

**Fig. 5.**
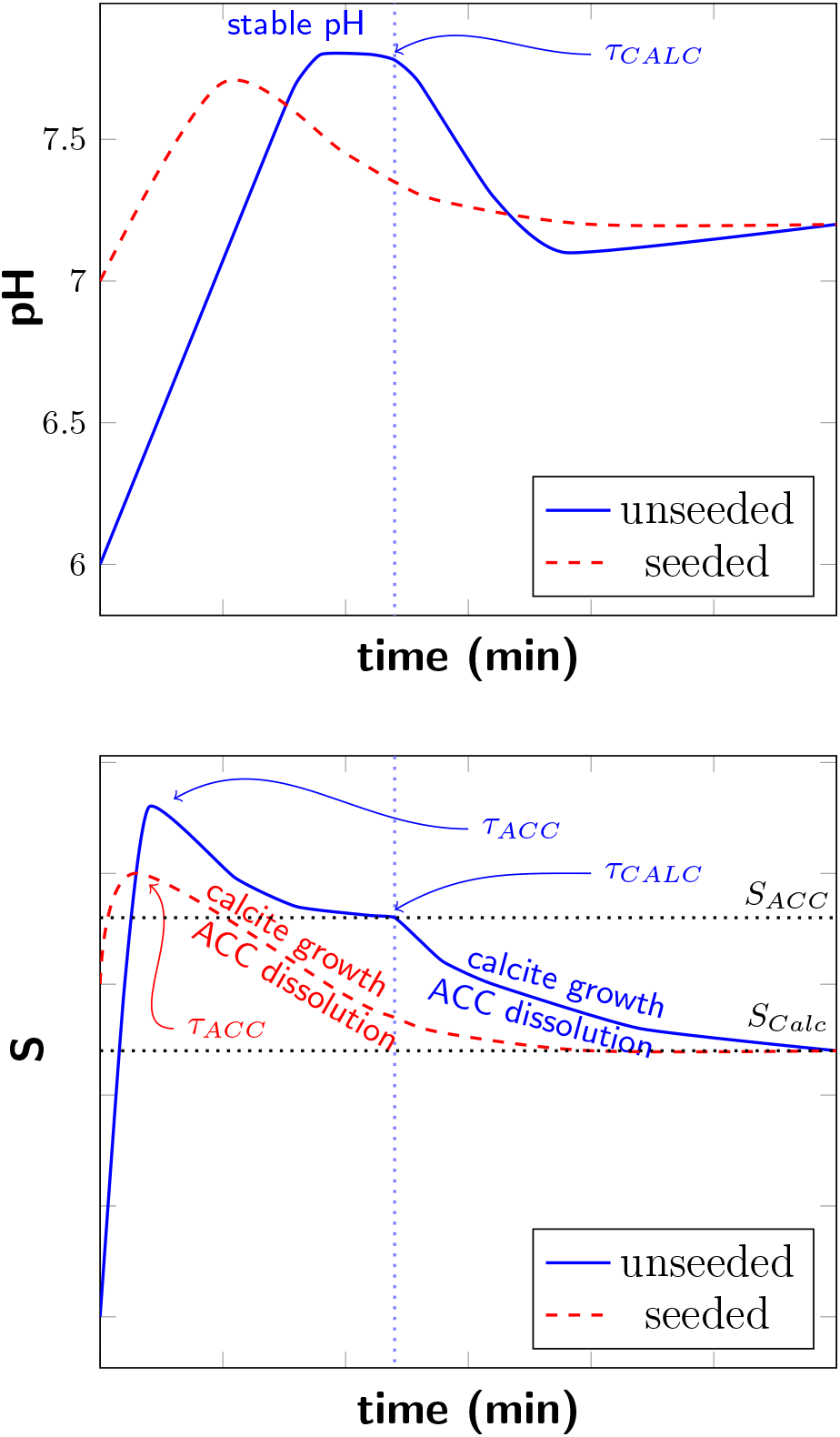
Schematic illustration of real time pH evolution (top) and saturation level (bottom) of unseeded and seeded samples with high bacteria cell concentration. *τ_ACC_* and *τ_CALC_* mark the nucleation of ACC and calcite. *S_ACC_* and *S_CALC_* represent *S* = 1 with respect to ACC and calcite, respectively.

In the starting phase of the experiment a rapid pH increase was observed for both unseeded and seeded samples. With increasing pH also the saturation in the sample is increasing. The pH increase was correlated to the urea hydrolysis rate, which was controlled by the bacteria cell concentration^24^. The bacteria cells were washed before the experiments, therefore there was no free enzyme present in the bacteria dilutions. However, the used bacteria strain (DSM33) has always urease enzyme with high activity present^25,26^, located most likely in the cell membrane. Therefore, urea hydrolysis and pH increase started just after adding the bacteria to the reaction well despite the fact that the timescale of the experiment was not sufficiently long for the bacteria to produce urease as a response to the presence of urea.

For unseeded samples, ACC nucleated (*τ_acc_*) once the saturation *S* was sufficiently above *S_ACC_* (*S_ACC_* represent *S* = 1 with respect to ACC) and the precipitation of ACC reduced *S*. Precipitated ACC transformed rapidly to vaterite according to Ostwald’s rule of stages (for simplification, this transformation is not shown in Figure 5). When the stable pH was reached, the urea hydrolysis rate by bacterial urease was equal to the precipitation rate. In this stable pH region the saturation level *S* is reduced to *S_ACC_* due to the growth of the ACC phase and stays constant until the nucleation of calcite occurs as schematically illustrated in Figure 5. Once calcite nucleated, nucleation and crystal growth of calcite contributed to reduce *S* and resulted in a pH decrease. The solution became under-saturated with respect to ACC/vaterite and the metastable phases dissolved.

With decreasing bacteria cell concentration, and therefore decreasing urea hydrolysis rate, a lower maximum pH values and correspondingly lower *S* were reached in the starting phase of the reaction. For lower *S* it takes longer until calcite nucleates and consequently a longer stable pH region was observed before the pH decreased (Figure 1a).

For the seeded samples a short stable pH region was observed for the three highest bacterial cell concentrations, and the shape of the pH curves was similar for those samples. For the lowest bacterial cell concentration no pH decrease was observed, and the pH stabilized after the initial pH increase (Figure 2a). The difference in the pH evolution for seeded samples could be connected to the precipitation of ACC and calcite. For the three highest bacteria cell concentrations, the pH increased until a short stable pH phase was reached, where the urea hydrolysis rate was equal to the precipitation rate, before the pH decreased. The pH decrease indicated that nucleation and growth on the calcite seeds was not sufficient to consume hydrolysis products. High pH and corresponding high supersaturation resulted in precipitation of ACC also for seeded samples. This was also confirmed by optical microscopy (Figure 2c). The pH decrease was a consequence of nucleation of calcite crystals and crystal growth. The initial calcite seeds acted as nucleation sites, and therefore no significant pH plateau was observed before calcite nucleation for the three highest bacterial cell concentrations. With decreasing pH and therefore decreasing *S* ACC/vaterite dissolved (see Figure 5, red dashed line). For the lowest bacterial cell concentration no pH decrease was observed. Due to a lower urea hydrolysis rate, caused by a lower bacteria cell concentration a lower saturation level was reached. In this case, the supersaturation produced by urea hydrolysis was most likely reduced by growth of existing calcite seeds and no ACC precipitated for the seeded sample with the lowest bacterial cell concentration. ACC/vaterite precipitation was reported earlier for unseeded samples^20,27^. However, we were able to show by comparing unseeded to seeded experiments that the metastable polymorph phases can also precipitate for samples where calcite seeds were present (Figure 2). This could also be confirmed by analyzing the shape of the pH evolution curves. The presence of calcite seeds lowered the energy barrier for nucleation, resulting that lower pH values were necessary for calcite nucleation. However, the presence of calcite seeds did not suppress the formation of metastable phases for higher bacteria cell concentrations.

Calcite nucleation and growth on the seeds started at lower pH values in seeded samples (Figure 2d) than calcite nucleation in unseeded samples and therefore at lower supersaturation. The influence of calcite seeds on CaCO_3_ precipitation in non-MICP experiments was reported earlier, and it was shown, that in the presence of calcite seeds, CaCO_3_ nucleation can take place at lower supersaturation levels^14,28^. In our experiments, we could confirm this also for MICP experiments. Furthermore, the calcite seeds had a significant influence on the precipitation process. With SEM characterization (Figure 4e-h) we could show that the seeds were connected together with precipitated CaCO_3_.

Furthermore, it was observed for unseeded samples that the bacterial cell concentration had a strong influence on the size and shape of the precipitated calcite crystals. It has been reported previously that a high supersaturation condition for nucleation leads to smaller CaCO_3_ crystal sizes ^29^. Cheng *et al*. and Cuberth *et al*. reported that the initial ureolysis rate in MICP experiments influenced the crystal size^30,31^. Wang *et al*. showed recently that higher bacteria densities in MICP experiments resulted in a high nucleation rate, due to the precipitation of unstable phases of CaCO_3_^24^. This was also observed in our experiments (Figure 1d and Figure 4a,c). A higher ureolysis rate in the beginning of the experiment resulted in a larger amount of ACC/vaterite precipitation, which could act as nucleation sites for calcite crystals. This resulted in the nucleation of a larger number of smaller crystals.

## 5 Conclusions

We presented a systematic study investigating the influence of calcite seeds on the crystallization processes of CaCO_3_ in MICP. In our experiments we observed a significant difference in the pH evolution between unseeded and seeded experiments, caused by different precipitation processes of CaCO_3_. By microscope analysis and analyzing the shape of the real time pH evolution and OD_750nm_ we could detect the precipitation of the metastable phases of ACC/vaterite also for samples with calcite seeds present. Furthermore, we detected that lower pH values were necessary for calcite nucleation in seeded experiments. We showed that the precipitation of metastable precipitates in seeded samples is connected to the urea hydrolysis activity, and can therefore be controlled by the bacterial cell concentration.

The precipitation of metastable phases is important in biocementation processes as it might affect the homogeneity of the consolidated MICP material. ACC particles are small and have a lower density compared to calcite and could therefore be transported with flow through the granular medium. Small pores in the granular medium might be clogged with ACC and therefore influence the homogeneity of calcite precipitation. The results obtained in this study show that also in the presence of initial calcite crystals ACC can precipitate for high bacterial cell concentrations and that no ACC precipitates in samples with low bacteria cell concentrations. This needs to be considered in MICP protocols to avoid clogging of the granular medium to achieve knowledge-based improvement of MICP materials.

## Acknowledgements

This work was supported by the Research Council of Norway under project 269084/O70. Additional support was provided by Norwegian Micro- and Nano-Fabrication Facility, NorFab under Research Council of Norway project 245963/F50. The authors thank Simone Balzer Le from SINTEF Industry, Trondheim, Norway, for providing the bacteria cultures for the performed experiments and Seniz Ucar from the department of Chemical Engineering, NTNU, Trondheim, Norway for the fabrication of the calcite seeds.

