## Supplementary document for "Calcite seed-assisted microbial induced carbonate precipitation (MICP) and its potential in biocementation"

### Methods

#### Scanning electron microscopy

The shape and the size of the fabricated calcite seeds was investigated with a scanning electron microscope (FEI APREO) (SEM). The dried calcite seeds were attached to a SEM stub with carbon tape and sputter coated (Cressington 208HR) with 10 nm platinum/palladium (80/20). SEM images were taken with an accelerating voltage of 2 kV.

### Supplementary Data

Table S1: Dilution of original bacteria cultures. The original culture with growth medium was centrifuged, washed and re-suspended in 0.01 M PBS. The re-suspended bacteria cultures without growth medium were dilute to the final dilutions, which were used for the experiments.

| | **Culture with growth medium** | | **Culture without growth medium** | | |
|---|---|---|---|---|---|
| **Sample name** | **Amount culture** | **PBS** | **Amount culture** | **PBS** | $\mathbf{OD_{600nm}}$ |
| D1-C1 | 4 mL | 4 mL | 1000 µL | 0 µL | 1.40 |
| D2-C1 | 4 mL | 4 mL | 750 µL | 250 µL | 1.30 |
| D3-C1 | 4 mL | 4 mL | 500 µL | 500 µL | 1.11 |
| D4-C1 | 4 mL | 4 mL | 200 µL | 800 µL | 0.64 |
| D1-C2 | 1 mL | 1.5 mL | 1000 µL | 0 µL | 0.96 |
| D2-C2 | 3 mL | 3.5 mL | 1000 µL | 0 µL | 0.88 |
| D1-C3 | 3 mL | 3.5 mL | 1000 µL | 0 µL | 0.89 |

Table S2: Starting pH value of the crystallization solution for samples with and without calcite seeds present before adding the bacteria culture to the crystallization solution.

| **Bacteria sample** | **pH unseeded** | **pH seeded** |
|---|---|---|
| D1-C1 | 5.3 | 6.1 |
| D2-C1 | 5.3 | 6.1 |
| D3-C1 | 5.3 | 6.3 |
| D4-C1 | 5.4 | 6.1 |

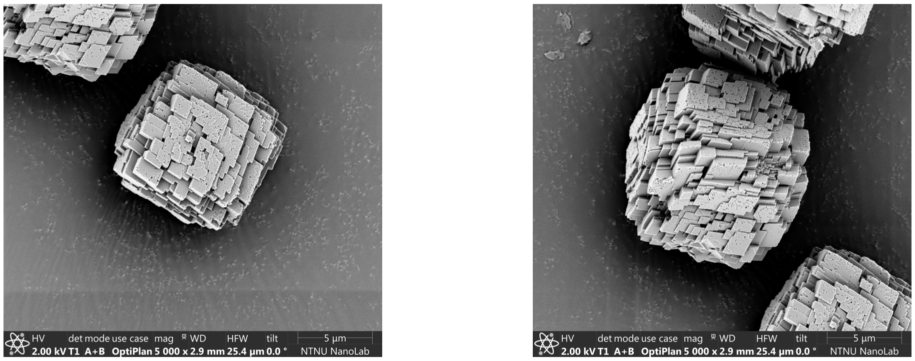


Figure S1: SEM images of calcite seeds.

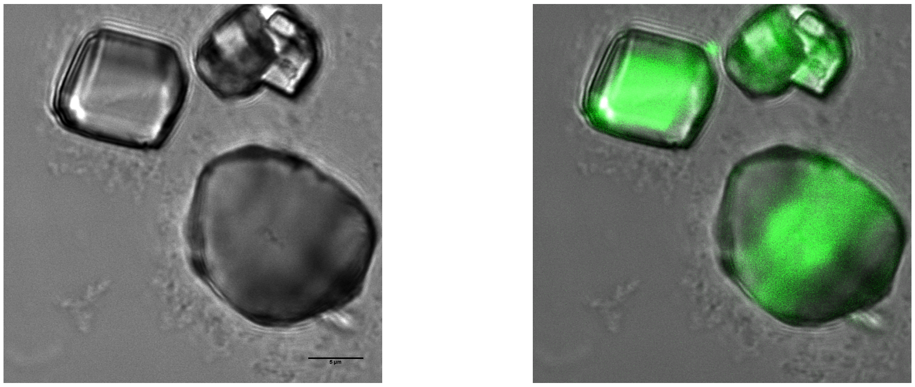

Figure S2: Confocal laser scanning microscope images of fluorescent calcite seeds grown on glass cover-slides. (left) Brightfield images of calcite seeds and (right) combination of brightfield image and fluorescent signal of incorporated fluorescent dye.
